# Distillation enables scalable high-fidelity virtual screening across ultra-large chemical libraries

**DOI:** 10.64898/2026.06.29.735361

**Authors:** Jiawei Dai, Yueyue Wang, Naing Lin Shan, Marco Mariani, Zimeng Yu, Qin Yan, Lalit K. Golani, Yulia V. Surovtseva, William H. Lee, Lajos Pusztai

## Abstract

Accurate virtual screening of ultra-large chemical libraries remains challenging. Existing approaches rely on lower-fidelity scoring functions or sampling-based strategies that can limit predictive accuracy and bias the exploration of chemical space. Here, we present FastBindRank, a distillation-based framework that transfers the predictive power of the structure-based model Boltz-2 into an efficient sequence-based surrogate. Trained on ∼1% of the 122-million-compound PubChem library, FastBindRank enables high-fidelity screening at scale. Applied to histone deacetylase 11 (HDAC11), FastBindRank substantially enriched high-confidence binders relative to the background chemical space. The lightweight model captured structural patterns associated with predicted binding, revealing structural determinants of binding. Under a comparable computational budget, FastBindRank achieved a 74-fold increase in hit rate and over a 30-fold increase in discovery yield over direct subset-based screening. Experimental validation confirmed the activity of two novel compounds. These results establish distillation as a practical strategy for scalable, high-fidelity virtual screening of ultra-large chemical libraries.

## Main

Identifying small molecules that bind to therapeutic targets is a central challenge in drug discovery. Computational screening has reshaped early-stage discovery by enabling rapid prioritization of large chemical libraries. Advances in structure-based modeling, including molecular docking methods (e.g. AutoDock^1^ and AutoDock Vina^2^), as well as deep learning-based approaches such as convolutional and graph neural networks^3, 4, 5^, and meta-learning frameworks incorporating docking-derived information^6^, have substantially improved prediction accuracy. More recently, models such as Boltz-2 have approached the performance of physics-based free-energy perturbation (FEP) methods in estimating protein–ligand binding affinity^7, 8^.

Despite these advances, a fundamental trade-off persists between predictive accuracy and scalability. High-accuracy structure-based models are computationally intensive and remain impractical for ultra-large library screening, often requiring substantial GPU resources^9^. As a result, virtual screening workflows are typically constrained to relatively small subsets of chemical space. While prior research has highlighted the importance of expanding chemical space in early-stage screening^10^, several strategies have been proposed, including machine learning-guided docking pipelines^11, 12^ and active learning-based approaches^13, 14, 15^. These methods share a common strategy of training models on compound subsets to guide large-scale prioritization. However, they typically rely on lower-fidelity proxies of binding affinity, such as docking scores, which can limit predictive accuracy. In addition, active learning strategies may be sensitive to sampling bias and model uncertainty, potentially prioritizing outliers or poorly represented regions of chemical space that provide limited value for identifying high-affinity binders.

We hypothesize that combining subset-based training with high-fidelity supervision can enable scalable screening without sacrificing predictive accuracy. Knowledge distillation, originally proposed to transfer predictive knowledge from a high-capacity teacher model to a more efficient student model, has become a widely adopted strategy for improving model efficiency while preserving predictive fidelity^16, 17, 18^.

Here, we apply target-specific knowledge distillation to transfer high-fidelity predictions from the computationally expensive structure-based model Boltz-2 to our novel deep learning model, BindRankNet, a lightweight compound sequence-based surrogate within the FastBindRank framework, for large-scale compound prioritization. We demonstrate the application of this novel framework by ranking the entire 122-million-compound PubChem library^19, 20^ to identify binders and potential inhibitors of histone deacetylase 11 (HDAC11). HDAC11, the sole class IV member of the histone deacetylase family, is an epigenetic regulator implicated in tumor progression and therapy resistance, but yet remains chemically underexplored with relatively few inhibitors available^21, 22, 23^. Two selected novel compounds were experimentally validated using an in vitro HDAC11 enzyme activity assay. In addition, we found that the lightweight model captured structural patterns of compounds associated with predicted binding, providing insight into structural determinants of ligand binding for targets. Our framework represents a practical and scalable method for high-fidelity virtual screening across ultra-large chemical spaces.

## Results

### Distillation-based design and performance evaluation of FastBindRank

We developed FastBindRank to prioritize protein-binding compounds at scale across the 122-million-compound PubChem library. As a case study, we applied this framework to HDAC11, an emerging epigenetic enzyme with a defined catalytic domain but a limited set of inhibitors with available experimental binding affinities (n = 8) (**Supplementary Fig. 1a** and **Supplementary Table 1**).

To select an optimal teacher model for structure-based distillation, we benchmarked multiple structure-based modes against this curated set of HDAC11 inhibitors (**Supplementary Table 2**). While AutoDock Vina^2^ showed no correlation with experimental pIC50 values (Pearson r = 0.031, *P* = 0.938) and GIGN^5^ and Boltzina^24^ exhibited only modest correlations (r = 0.332 and 0.368, respectively; *P* = 0.319 and 0.296, respectively), Boltz-2^8^ showed a strong statistically significant correlation (r = 0.815, *P* = 0.002) (**Supplementary Fig. 1b–e**). Moreover, Boltz-2 predicted pIC50 values correlated positively with its binding probability estimates (r = 0.637, *P* = 0.035; **Supplementary Fig. 1f**), supporting its selection as the teacher model for distillation.

The student ranking model, BindRankNet, was trained to estimate Boltz-2 binding probabilities using Morgan fingerprints of compounds, which provide one-dimensional representations of molecular structure (**Fig. 1**). A randomly sampled set of 1.85 million compounds was partitioned into independent training (n = 250,000 per iteration), tuning (n = 100,000), and two independent test splits (n = 250,000 each). Across five independent training iterations, this design required Boltz-2 predictions for only ∼1% of the 122-million-compound PubChem library, enabling efficient model development while preserving representative chemical diversity (**Fig. 1a**).

**Fig. 1:**
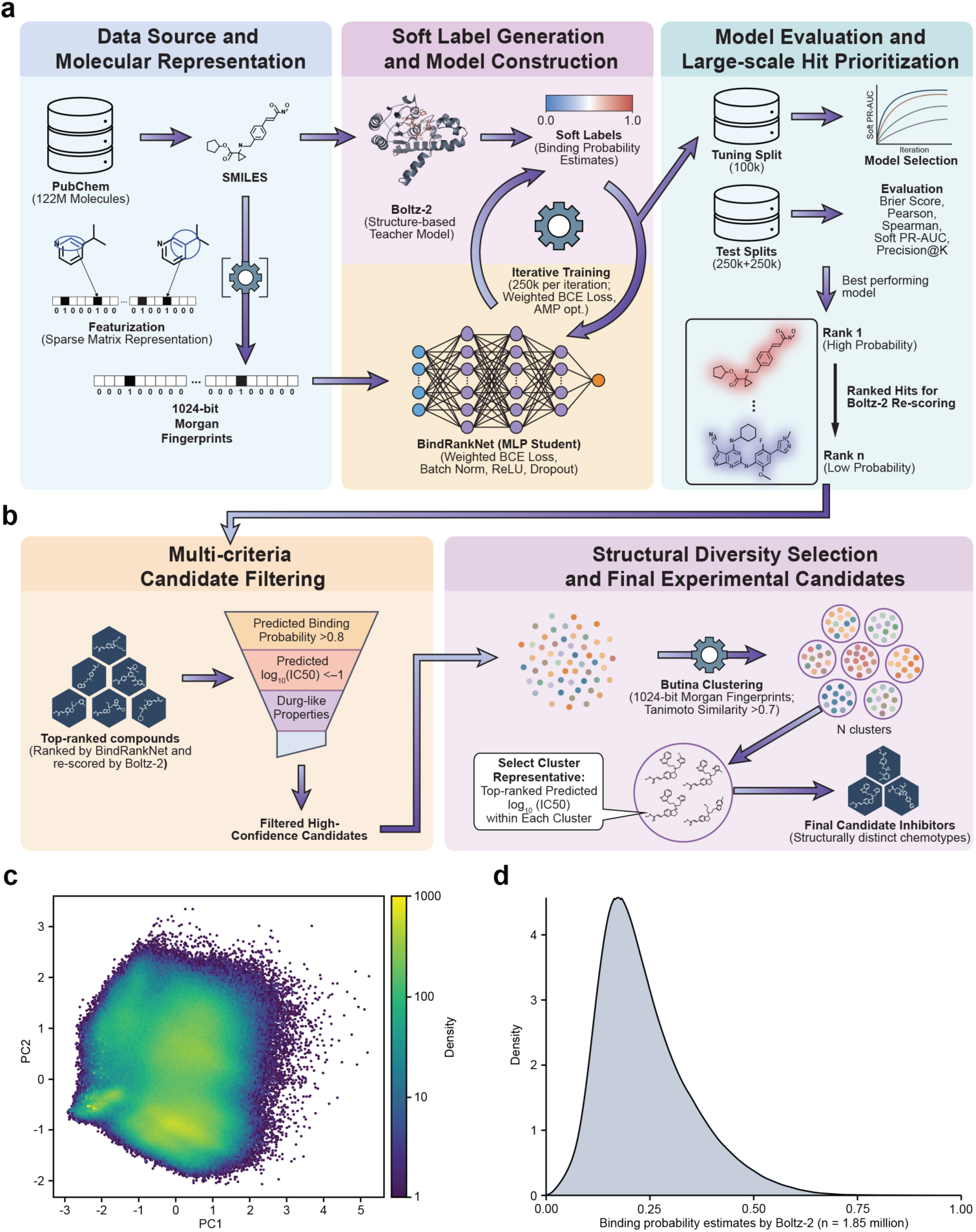
FastBindRank workflow and candidate selection. **a,** PubChem compounds are represented using molecular fingerprints. Boltz-2 predictions are used to train BindRankNet, enabling model evaluation and large-scale score-based hit prioritization. **b,** Top-ranked compounds are filtered by predicted binding strength and drug-like properties, followed by structural diversity selection to obtain final candidate inhibitors. **c,** Chemical space of the 1.85 million compounds used for model training, tuning, and testing in the HDAC11 case study. **d,** Binding probability distribution predicted by Boltz-2 for HDAC11 across the same compound set as in panel c.

Systematic hyperparameter optimization identified a configuration that maximized the soft Precision–Recall Area Under the Curve (PR-AUC) on the tuning split (**Fig. 2a**). The final model demonstrated strong performance on the training data aggregated across five iterations (**Fig. 2b**) and generalized well to the tuning split, both independent test splits, and a curated set of known HDAC11 inhibitors (**Fig. 2c–f**). Concordance analyses further revealed strong correlations and favorable calibration (low Brier scores) between BindRankNet predictions and the Boltz-2 teacher (**Fig. 2g–i**). Performance converged rapidly across iterations, indicating that high-fidelity structural supervision provides strong learning signals early in training (**Supplementary Fig. 2**).

**Fig. 2:**
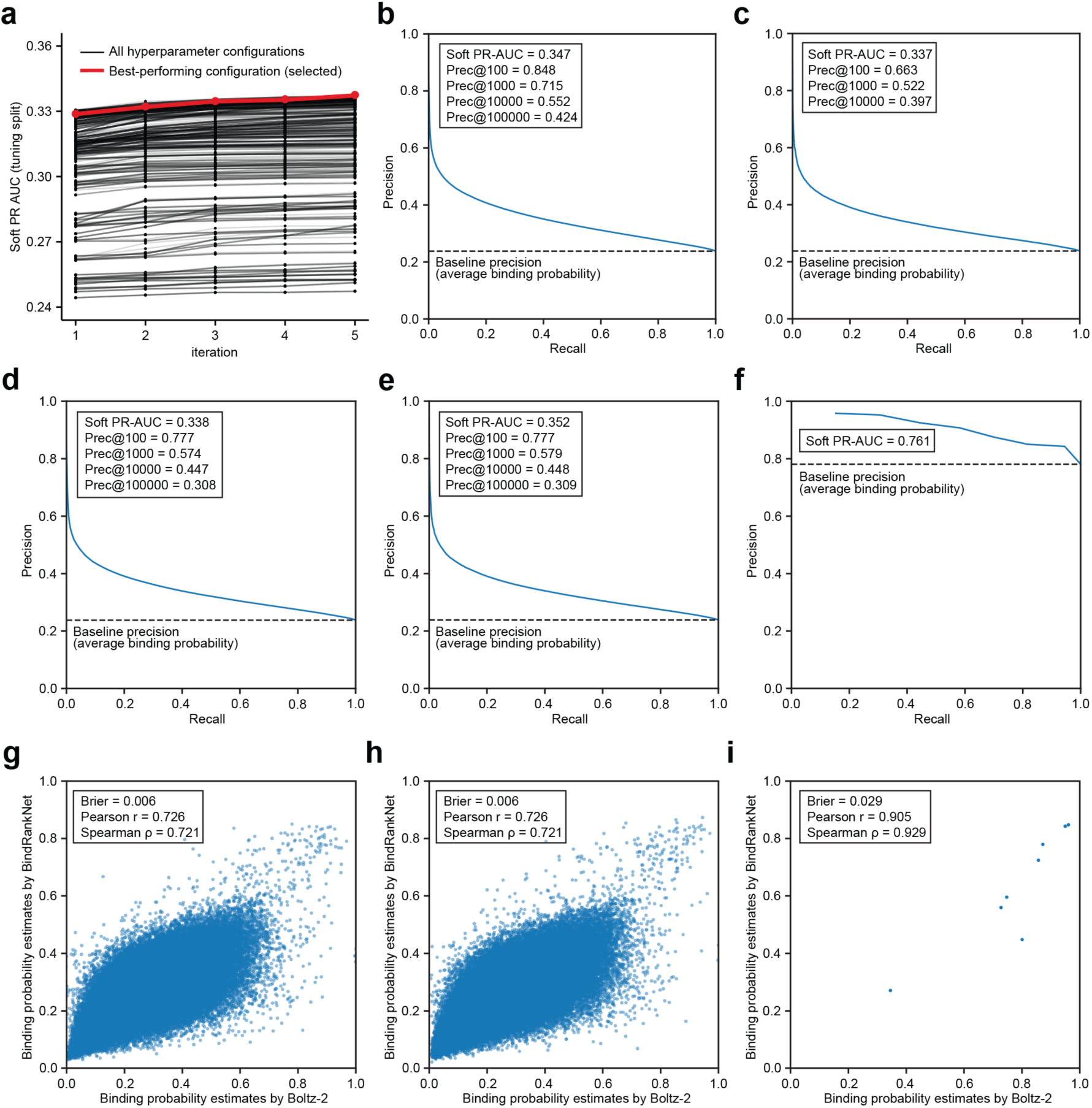
Performance evaluation of BindRankNet for structure-based prediction. **a,** Performance of BindRankNet across iterative training rounds during systematic hyperparameter search. Each black line represents one hyperparameter configuration, and the red line denotes the best-performing configuration selected based on soft PR-AUC on the tuning split. **b–f,** Soft Precision-Recall (PR) curves of the selected model evaluated on the training data aggregated across five independent iterations (**b**), tuning split (**c**), test split 1 (**d**), test split 2 (**e**), and a curated set of known HDAC11 inhibitors (**f**). Dashed lines indicate baseline precision, defined as the average binding probability in each dataset. Soft PR-AUC and Precision@K are shown. **g–i,** Concordance between binding probability estimates predicted by BindRankNet and Boltz-2 for compounds on test split 1 (**g**), test split 2 (**h**), and a curated set of known HDAC11 inhibitors (**i**). Brier scores, Pearson correlation coefficients (r), and Spearman correlation coefficients (ρ) are shown.

Following model selection and early prioritization, the FastBindRank framework incorporates structure-based re-scoring (Boltz-2), multi-criteria filtering (binding probability, log₁₀(IC50) and drug-like molecular properties), and structural diversity selection via Butina clustering^25^ (**Fig. 1b**). Analysis of the 1.85-million-compound subset confirmed coverage of a broad region of chemical space (**Fig. 1c**). Crucially, the distribution of Boltz-2–predicted binding probabilities was highly right-skewed, with only a small fraction of compounds showing elevated binding potential (**Fig. 1d**). This intrinsic sparsity of high-confidence binders underscores the importance of large-scale ranking to efficiently identify promising candidates.

### Efficient ranking across the 122-million-compound PubChem library

We applied the optimized BindRankNet model to screen the entire 122-million-compound PubChem library. As expected, the predicted score distribution was highly skewed, with the majority of compounds showing low probability estimates and a small fraction forming a tail of higher-scoring candidates (**Fig. 3a**). This distribution is consistent with the sparse distribution of active compounds in chemical space.

**Fig. 3:**
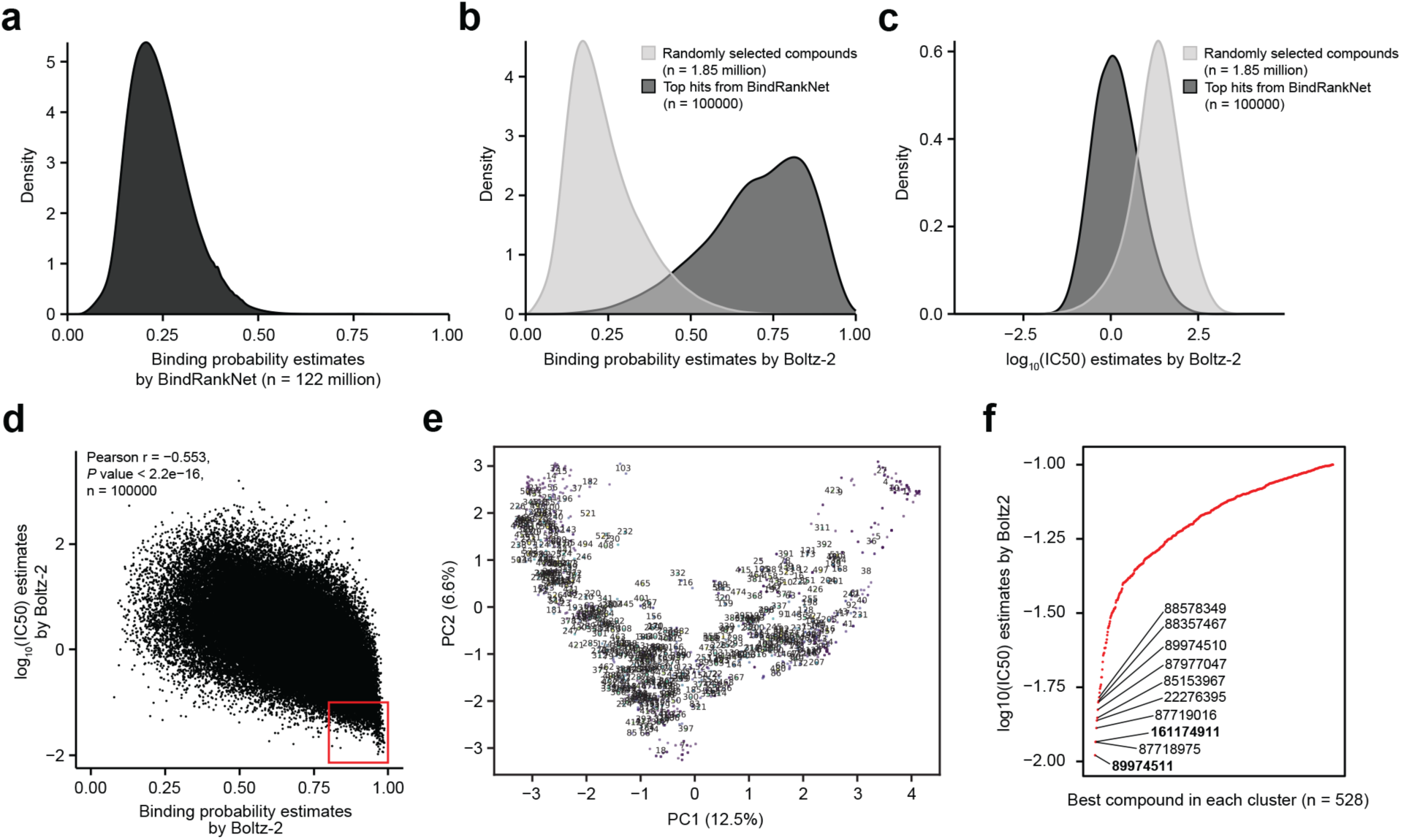
Large-scale virtual screening and candidate prioritization using FastBindRank. **a,** Distribution of binding probability estimates produced by BindRankNet for all 122 million PubChem compounds. **b,c,** Distribution of structure-based re-scoring results by Boltz-2 applied to FastBindRank top hits. Density plots show (**b**) Boltz-2–predicted binding probabilities and (**c**) Boltz-2–predicted log₁₀(IC50) values for the top 100,000 compounds prioritized by FastBindRank (dark grey), relative to a randomly sampled subset representing the background chemical space (light grey). **d,** Relationship between Boltz-2–predicted binding probabilities and log₁₀(IC50) estimates for BindRankNet top hits (n = 100,000) after re-scoring. Pearson correlation coefficient (r), corresponding *P* value, and number of compounds are shown. The red box highlights compounds with both high predicted binding probability (>0.8) and low predicted log_10_(IC50) (<−1). **e,** PCA-based visualization of the chemical space of BindRankNet top hits after filtering, illustrating structural diversity among prioritized compounds (n = 1,262). Numbers denote cluster identifiers. **f,** Rank-ordered Boltz-2–predicted log₁₀(IC50) values for representative compounds selected from each structural cluster (n = 528). The top 10 compounds with lowest log₁₀(IC50) values are labeled and the compounds selected for experimental validation are shown in bold.

To evaluate ranking performance, we compared the top 100,000 prioritized compounds against the full distribution represented by the randomly sampled 1.85-million-compound subset. Structure-based re-scoring showed that the BindRankNet-prioritized set was strongly enriched for higher Boltz-2 binding probabilities (Cliff’s δ = 0.97; **Fig. 3b**) and more potent predicted log₁₀(IC50) values (Cliff’s δ = −0.75; **Fig. 3c**) (two-sided Wilcoxon rank-sum test, *P* < 2.2 × 10⁻¹⁶ for both). These results indicate that FastBindRank effectively concentrates high-confidence binders at the top of the ranking despite training on only ∼1% of the library.

We then applied joint thresholds of binding probability > 0.8 and log₁₀(IC50) < −1 to the top-ranked set, reducing the pool to 2,043 compounds (**Fig. 3d**). Subsequent filtering for drug-like molecular properties yielded 1,262 high-confidence candidates (**Supplementary Table 3**). This selection process recovered the known HDAC11 inhibitors JNJ-26481585 and Panobinostat, which are the strongest known experimentally validated inhibitors, providing support for the validity of our screening approach (**Supplementary Table 1**). Most importantly, many newly identified candidates were predicted to have comparable or greater inhibitory potency than these known inhibitors.

### Identification of structurally diverse and novel candidate compounds

To ensure structural diversity among these high-confidence candidates, we performed Butina clustering based on Morgan fingerprints. Principal component analysis (PCA)^26^ confirmed that the prioritized candidates span a broad chemical structure space (**Fig. 3e**). The clustering process identified 528 distinct clusters, characterized by a maximum cluster size of 55 and a molecule-weighted median size of 4, indicating substantial structural diversity within the candidate pool. From each cluster, we selected the representative compound with the highest Boltz-2–predicted binding affinity, yielding a focused set of 528 structurally diverse candidates. Importantly, selecting one representative per cluster retained high predicted binding affinity while enforcing structural diversity (**Fig. 3f**).

Finally, to prioritize novelty, we cross-referenced the representatives with CAS SciFinder^27^ to exclude annotated histone deacetylase (HDAC) binders. This filtering step reduced the set to 454 novel candidate compounds with no prior HDAC annotation. Detailed information for these candidates is provided in **Supplementary Table 3**.

### Structural patterns associated with predicted binding

To interpret the structural patterns underlying model predictions, SHapley Additive exPlanations (SHAP)^28^ analysis was performed on the final BindRankNet model. Top-ranked candidates (n = 100,000) were used to identify Morgan fingerprint bits that contributed most strongly to predicted binding probability (**Supplementary Table 4**). Global SHAP analysis revealed a subset of fingerprint bits with high contributions, indicating that model predictions are driven by specific local chemical environments (**Supplementary Fig. 3a**).

Representative atom environments corresponding to the high-SHAP fingerprint bits were identified across multiple compounds selected based on the highest SHAP values for each bit, revealing recurring local structural patterns associated with predicted HDAC11 binding (**Supplementary Fig. 3b**). At the molecular level, representative candidate compounds showed that top-ranked candidates frequently contain multiple high-SHAP-associated substructures (**Supplementary Fig. 3c**). To evaluate the robustness of these feature importance patterns before and after filtering, SHAP values computed for the top-ranked candidates (n = 100,000) and final representative candidates (n = 528) were compared. A strong concordance was observed between the two sets (Pearson r = 0.990, *P* < 2.2 × 10⁻¹⁶) (**Supplementary Fig. 3d**), demonstrating that the structural features learned by the model are largely preserved following physicochemical filtering and structural diversity selection. Together, these results suggest that the model captures structural patterns that are reflected in the local chemical environments of predicted HDAC11 binders and are consistent across candidate selection stages.

### Benchmark comparison under matched computational budget

Directly applying Boltz-2 to the full 122-million-compound PubChem library is computationally prohibitive, requiring an estimated 91,811 GPU days (**Supplementary Table 5**). To establish a feasible structure-based baseline, we performed direct Boltz-2 screening on a randomly sampled subset of 1.85 million compounds, corresponding to the same subset used for model training and evaluation, and serving as a computationally matched reference for comparison.

Within this subset, BindRankNet predictions showed strong correlation with Boltz-2 scores (Pearson r = 0.764; Spearman ρ = 0.757; **Supplementary Fig. 4a**), indicating that the model captures Boltz-2–derived binding signals. Applying identical filtering criteria, we found only 17 high-confidence candidates, which were further reduced to 16 structurally representative compounds after clustering (**Table 1**, **Supplementary Fig. 4b–d and Supplementary Table 6**).

**Table 1.** Quantitative benchmark of compound prioritization outcomes under comparable computational budgets.

| Metric | Screening strategy |  |  |
| --- | --- | --- | --- |
|  | FastBindRank (full PubChem; top 100,000) | Boltz-2 (1.85 million subset; top 100,000) | Boltz-2 (1.85 million subset; proportional selection) |
| Total compounds screened | 122 million | 1.85 million | 1.85 million |
| Top hits selected | 100,000 (0.082%) | 100,000 | 1,516 (0.082%) |
| Compounds passing filters (pre-diversity selection) | 1,262 | 17 | 17 |
| Average Boltz-2 binding probability (filtered) | 0.919 (0.801 to 0.994) | 0.898 (0.804 to 0.970) | 0.898 (0.804 to 0.970) |
| Average Boltz-2 $\log_{10}(\text{IC}_{50})$ (filtered) | -1.19 (-1.98 to -1.00) | -1.24 (-1.59 to -1.04) | -1.24 (-1.59 to -1.04) |
| Clusters after diversity selection | 528 | 16 | 16 |
| Estimated wall-clock time (days) | 1,473.53 | 1,387.48 | 1,387.48 |

Selecting 100,000 compounds from the full PubChem library corresponds to approximately 0.082% of the total library; applying the same proportion to the 1.85M subset yields 1,516 top-ranked compounds. The filtering criteria applied before structural diversity selection include binding probability $> 0.8$ , $\log_{10}(\text{IC}_{50}) < -1$ , and drug-like molecular properties. Wall-clock time was measured or estimated using a single NVIDIA GPU.

Under matched computational cost, FastBindRank markedly improved discovery efficiency. While direct subset screening identified only 17 high-confidence candidates, FastBindRank identified 1,262 high-confidence candidates from the full library using a comparable computational budget (1,474 vs. 1,387 GPU days), representing a 74-fold increase in hit rate. This set was further refined to 528 structurally diverse representatives, corresponding to over a 30-fold increase in discovery yield relative to direct subset screening (**Table 1**).

To further compare prioritization behavior, we examined the overlap between compounds selected by BindRankNet and those identified by Boltz-2 within the subset (**Fig. 4a**). Most compounds selected by Boltz-2 within the subset (Venn region A; 97.8%) exhibited weak predicted binding potential. In contrast, BindRankNet prioritized compounds in Venn regions D and E, which were enriched for higher predicted binding probabilities and lower predicted log₁₀(IC50) values (**Fig. 4b–c**). Within a proportional selection set (n = 1,516), compounds jointly prioritized by both approaches (region D) consistently showed stronger predicted binding than those uniquely identified by Boltz-2 (region B).

**Fig. 4:**
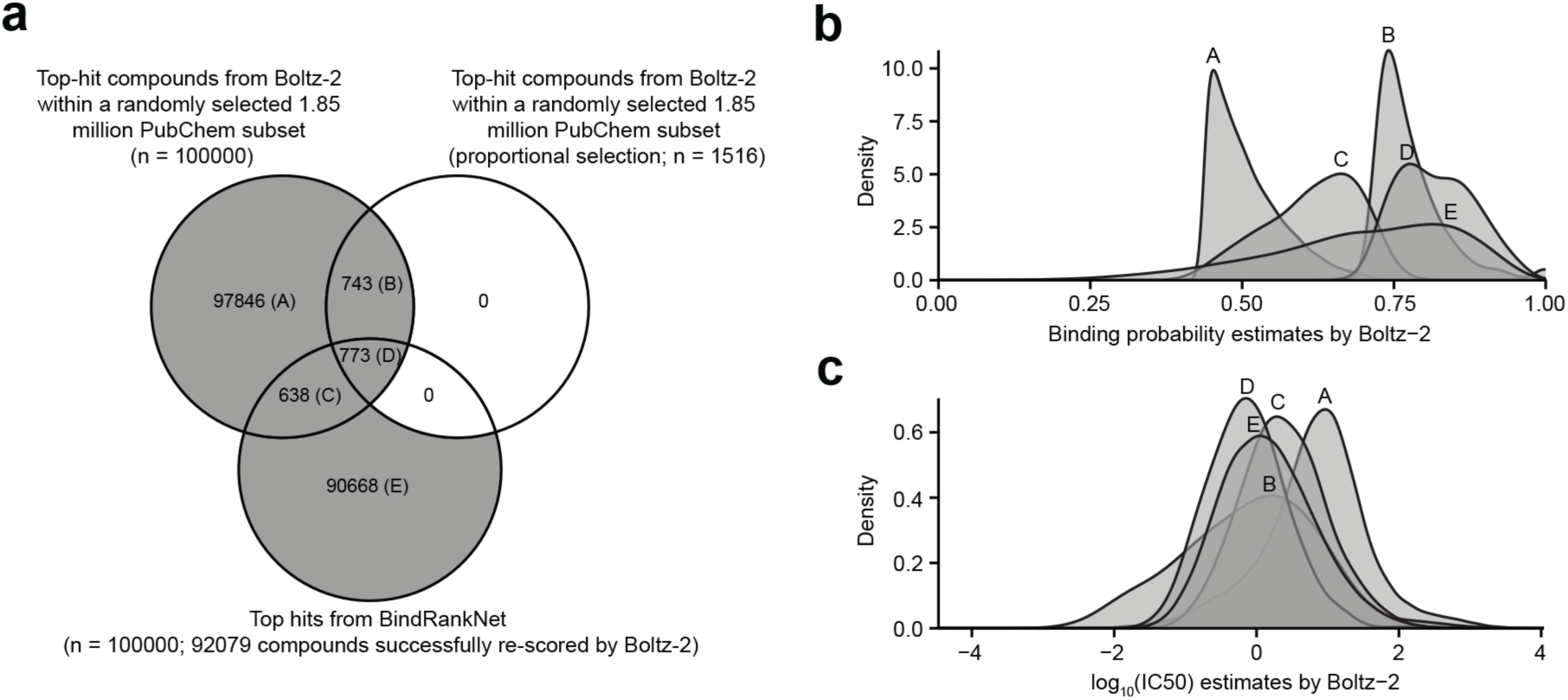
Benchmark comparison of prioritization performance under matched computational budget. **a,** Overlap between compounds prioritized by BindRankNet and those identified by direct Boltz- 2 screening within a randomly sampled subset of 1.85 million PubChem compounds. **b,c,** Distributions of Boltz-2–predicted (**b**) binding probabilities and (**c**) log₁₀(IC50) values for compounds in the Venn regions shown in panel a.

BindRankNet did not recover all high-affinity candidates identified by direct structure-based screening, as reflected by a small subset of compounds in region B. In addition, BindRankNet selected a minor fraction of compounds with relatively weak predicted binding potential (region C). However, because structure-based re-scoring is restricted to a small fraction of top-ranked compounds, inclusion of these cases has minimal impact on overall computational efficiency.

### Experimental validation of two novel HDAC11 inhibitor lead compounds

To experimentally evaluate prioritized candidates, we selected two compounds (PubChem CIDs 89974511 and 161174911) from the 454 novel candidates based on their high predicted binding affinity, low predicted log₁₀(IC50) values, predicted ease of chemical synthesis, and costs. The final two compounds were custom synthesized and tested for HDAC11 inhibitory activity in an in vitro enzyme activity assay (**Fig. 5a**). We selected Panobinostat and Fimepinostat as positive controls from our curated set of known HDAC11 inhibitors. These were selected because they represent two US Food and Drug Administration approved non-specific HDAC inhibitors used in the clinic to treat hematological malignancies.

**Fig. 5:**
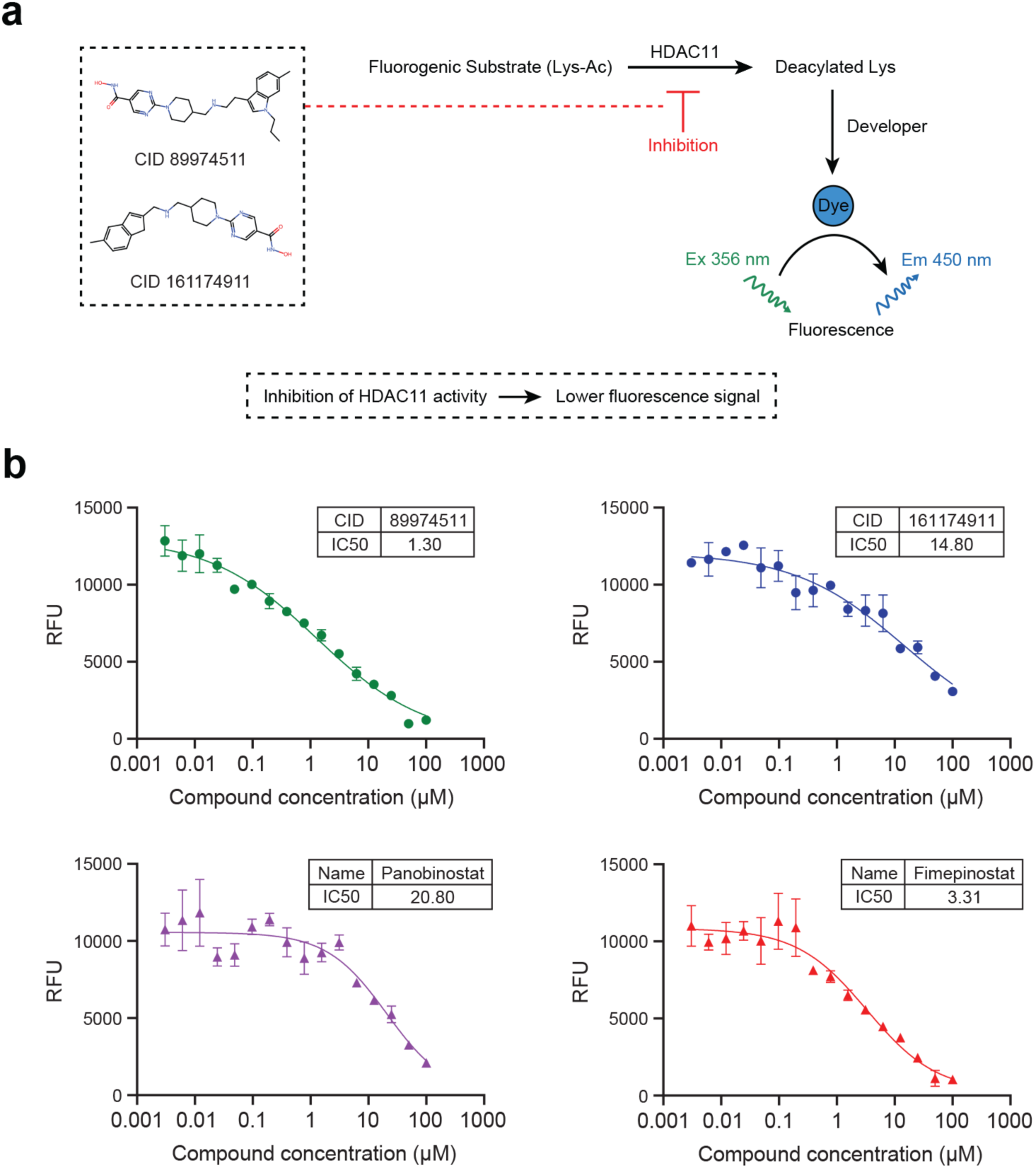
Inhibition of HDAC11 by selected compounds. **a,** Selected compounds (PubChem CIDs 89974511 and 161174911) evaluated for inhibition of HDAC11 using a fluorogenic assay, in which enzyme activity is reflected by fluorescence output. Inhibitors reduce fluorescence signal. **b,** Dose–response curves and IC₅₀ values for the two test compounds and positive control inhibitors (Panobinostat and Fimepinostat). RFU, relative fluorescence units.

Both novel compounds showed HDAC11 inhibitory activity similar to or higher than the positive controls. The IC50 values of CID 89974511 and CID 161174911 were 1.3 µM and 14.8 µM, respectively, which compare favorably with Panobinostat and Fimepinostat that had IC50 values of 20.8 µM and 3.3 µM, respectively (**Fig. 5b**). In our assay, CID 89974511 showed potency exceeding both clinically approved HDAC inhibitors. These results provide experimental support for the predictive capability of FastBindRank and demonstrate that large-scale structure-guided prioritization can identify functionally active novel compounds, offering a practical path for candidate discovery from ultra-large virtual screening.

## Discussion

In this study, we demonstrate that distillation can be effectively applied to bridge the long-standing gap between predictive accuracy and scalability in structure-based virtual screening. By transferring high-fidelity predictions from a computationally intensive structure-based model to a lightweight sequence-based surrogate, FastBindRank enables efficient prioritization across ultra-large chemical libraries such as PubChem while preserving key predictive characteristics of the teacher model. Although the teacher model’s predictions are not ground truth, this study shows that using high-fidelity labels is a reasonable strategy to build a scalable surrogate model with high accuracy. In addition, our previous work showed that sequence-based deep learning models were competitive to structure-based deep learning models in ligand-protein binding affinity prediction^6^, supporting the scalability and high performance of FastBindRank, particularly given the substantially greater computational efficiency of sequence-based models. We also note that the target-specific knowledge distillation does not require protein target information as input in BindRankNet, further improving efficiency of the student model. In the HDAC11 case study, although the model was trained using only ∼1% of the full chemical library, it retained strong agreement with Boltz-2 and supported effective large-scale prioritization. This approach yielded a 74-fold increase in hit rate and over a 30-fold increase in discovery yield under a matched computational budget, highlighting its practical advantage for large-scale screening. Rapid performance convergence across iterations further suggests that high-fidelity structural supervision provides strong early learning signals, enabling effective models to be trained with less data. Reproducible structural patterns associated with target binding are captured by the model, suggesting the potential to explore structural determinants of ligand binding for targets.

A major challenge in large-scale virtual screening is the intrinsic sparsity of high-affinity binders within chemical space, a “needle-in-a-haystack” problem^29, 30^. This is illustrated by our result that Boltz-2–predicted binding probabilities are highly skewed, with only a small fraction of compounds exhibiting high binding potential. This feature is the primary driver of the prohibitively high computational cost of directly applying high-fidelity structure-based models, whereas random or subset-based screening remains inefficient. This is reflected in the limited number of high-confidence candidates obtained from direct structure-based screening, compared with the substantially enriched candidates identified by FastBindRank.

Compared with previously proposed acceleration strategies, FastBindRank addresses a central limitation by directly leveraging high-fidelity structure-based supervision. Existing machine learning–guided docking pipelines^11, 12^ and active learning–based approaches^13, 14, 15^ aim to improve screening efficiency by selectively evaluating compounds, often through iterative or surrogate-guided selection strategies. In practice, these approaches rely on approximate signals, which can limit predictive accuracy, as well as sampling-driven selection rules that may introduce bias toward specific regions of chemical space. In contrast, FastBindRank applies distillation to transfer predictive behavior from a high-accuracy structure-based model to a scalable sequence-based surrogate, thereby reducing reliance on lower-fidelity proxy signals and improving predictive consistency at scale. This difference is particularly relevant given that docking-based scoring functions show limited agreement with experimental inhibitory activity, as observed in our analysis. Moreover, FastBindRank is trained on randomly sampled subsets that are independent of model predictions, rather than relying on model-driven iterative selection, thereby reducing sampling bias and providing a more uniform coverage of chemical space.

The identification of 1,262 high-confidence candidates and 528 structurally diverse representatives, together with the experimental validation of two compounds exhibiting inhibitory activity against HDAC11 that exceeds those of clinically used inhibitors, provides strong support for the practical utility of our approach. We show that candidates prioritized through FastBindRank have the predicted biochemical activity.

Several limitations should be acknowledged. First, FastBindRank is designed to approximate high-fidelity structure-based predictions rather than to directly predict experimentally measured inhibitory activity. The effectiveness of the framework therefore depends on the quality and generalizability of the teacher model. Second, the model is trained in a target- specific manner, requiring retraining for each new protein, and it remains unclear whether transfer learning from pre-trained models can be effectively applied across targets. Third, experimental validation was performed on only two carefully selected compounds due to cost limitations, and further studies will be required to systematically assess hit rates and optimize compound potency. Finally, the current implementation relies on ligand-based one-dimensional representations, which may not fully capture the three-dimensional geometry of protein–ligand interactions. In addition, the current framework does not generate explicit protein–ligand binding poses.

Despite these limitations, the results highlight a generalizable strategy for scaling high-fidelity structure-based predictions to ultra-large chemical libraries. As structure-based models continue to improve, distillation-based frameworks such as FastBindRank offer a practical route for extending these advances to large-scale discovery pipelines.

## Methods

### Catalytic domain of human HDAC11

The catalytic tunnel of human HDAC11 is mainly constituted by Gly151, Phe152, His183, Tyr209, Leu268, and Tyr304^31, 32^. The amino acid sequence within the catalytic domain was extracted for analysis (**Supplementary Fig. 1a**).

### Compound library curation and molecular representation

SMILES representations of 122,224,403 small-molecule compounds were retrieved from the PubChem database (https://pubchem.ncbi.nlm.nih.gov/), constituting a large and chemically diverse screening library^19, 20^. Molecular structures were transformed into 1024-bit Morgan fingerprints (radius = 2, with chirality information included) using RDKit (version 2024.3.5)^33^, providing a compact and standardized representation of local chemical environments and stereochemical features.

Morgan fingerprints were selected as a widely used and scalable molecular representation suitable for large-scale cheminformatics analyses. This representation was used consistently across downstream analyses, including chemical space visualization, binding probability modeling and prediction, as well as structural diversity assessment.

### Chemical space visualization of PubChem compounds

To visualize the global chemical space represented in PubChem, we randomly sampled 1.85 million compounds (1.5%) from the full library and projected their molecular fingerprints into a low-dimensional space using principal component analysis (PCA)^26^. An incremental PCA implementation was employed to enable efficient processing of the large dataset in mini-batches (batch size = 50,000) without requiring the full feature matrix to be loaded into memory. The trained PCA model was subsequently used to obtain two-dimensional embeddings for all sampled compounds, defined by the first two principal components (PC1 and PC2).

The distribution of compounds in PCA space was visualized using hexagonal binning to represent local compound density, which provides a robust summary for large-scale datasets. Color intensity corresponds to the number of compounds per bin and is displayed on a logarithmic (log₂) scale.

### Collection of known HDAC11 inhibitors and computational binding affinity prediction

Eight HDAC11 inhibitors with available experimental binding affinities were curated from the literature and public resources (**Supplementary Table 1**). Reported half-maximal inhibitory concentrations (IC50) from enzyme inhibition assays were converted to molar units and transformed to pIC50 (−log₁₀IC50) for comparison.

Binding affinity predictions for these inhibitors were generated using four computational tools: Boltz-2 (version 2.2.0), AutoDock Vina (version 1.2.7), GIGN and Boltzina (version 0.1.0)^2, 5, 8, 24^. Boltz-2 was run with its default parameters to predict HDAC11–ligand affinities. Multiple sequence alignments (MSAs) required for protein representation were generated using the built-in automatic MSA pipeline based on ColabFold^34^. For AutoDock Vina and Boltzina, the ligand binding site of HDAC11 was defined by P2Rank (version 2.5) using the AlphaFold DB structure of HDAC11 (UniProt: AF-Q96DB2-F1), obtained via UniProt (https://www.uniprot.org/)^35, 36, 37^. The docking grid box was set to 20 × 20 × 20 Å³, with exhaustiveness = 32. For GIGN, ligand–protein binding poses were first generated by Chai-1 (version 0.6.1) with default settings^38^ and the resulting complexes were then used as input to GIGN to estimate binding affinities. Boltzina was executed in full docking mode under default parameters.

### Model architecture and training

BindRankNet, a multilayer perceptron (MLP) model (student model) was developed to predict HDAC11 Boltz-2 (teacher model) binding probabilities for compounds using 1024-bit Morgan fingerprints from SMILES strings of compounds. The network comprised fully connected layers with batch normalization, rectified linear unit (ReLU) activation, and dropout regularization. Model training employed a weighted binary cross-entropy loss with logits (BCEWithLogitsLoss), and optimization was performed using adaptive moment estimation (Adam), Adam with weight decay regularization (AdamW), or stochastic gradient descent (SGD).

During training, the dataset was organized such that the tuning split remained fixed across all iterations, while the training split was randomly resampled in each iteration. Boltz-2 was used to predict the binding probability of each molecule toward HDAC11 as the soft label for BindRankNet. In each iteration, 250,000 compounds were used for training, and early stopping based on loss convergence was applied to determine training termination. Specifically, when the loss on the tuning split did not decrease for three consecutive epochs, the model corresponding to the lowest loss was retained as the final model for that iteration. Each iteration was initialized from the model of the previous iteration to enable progressive refinement of learned representations. Automatic mixed-precision (AMP) optimization was employed to improve computational efficiency and accelerate convergence during training.

Hyperparameters were systematically tuned across multiple configurations. Network architectures with four predefined configurations were evaluated: 1024–512–256, 2048–1024–512, 2048–1024–512–256, and 4096–2048–1024–512 neurons across successive hidden layers. Dropout regularization was applied with rates of 0.0, 0.1, 0.2, or 0.3. Learning rates of 1×10⁻⁶, 1×10⁻⁵, 1×10⁻⁴, and 1×10⁻³ were examined, and the L2 weight decay regularization coefficient was varied across 1×10⁻⁶, 1×10⁻⁵, 1×10⁻⁴, 1×10⁻³, and 1×10⁻² for AdamW and SGD optimizers. A momentum value of 0.9 was applied for SGD optimization only. All models were implemented in PyTorch (version 2.7.1) and trained on NVIDIA GPUs.

After five iterations, a fixed tuning split consisting of 100,000 randomly sampled compounds was used to assess performance variation across iterations and hyperparameter combinations. Model selection was based on soft Precision–Recall Area Under the Curve (PR-AUC), in which continuous probabilistic labels were used to compute precision and recall without binarization, thereby directly reflecting ranking quality for experimental prioritization. The selected best performing model was finally evaluated on two independent test splits, each comprising 250,000 randomly sampled compounds. Performance was assessed using the Brier score to evaluate probability calibration; Pearson (r) and Spearman (ρ) correlation coefficients to assess ranking consistency; soft PR-AUC to measure enrichment of likely binders toward the top of the ranked list; and Precision@K to quantify the average binding probability assigned by the teacher model among the top K ranked compounds.

### Large-scale virtual screening and candidate selection

The best-performing model was subsequently applied to predict HDAC11 binding probabilities for all 122,224,403 compounds in the PubChem library. The top 100,000 ranked compounds (0.082% of the library) were selected for Boltz-2 re-scoring. In parallel, molecular physicochemical properties for these top-ranked compounds (top hits) were deterministically computed from SMILES strings using RDKit, including molecular weight (MW), calculated octanol/water partition coefficient (cLogP), hydrogen bond donors (HBD), hydrogen bond acceptors (HBA), topological polar surface area (TPSA), number of rotatable bonds (RotB), total ring count (RingCount), number of aromatic rings (AromaticRings), number of heavy atoms (HeavyAtoms), and formal molecular charge (FormalCharge). Drug-like compounds were defined using the following criteria: MW 250–550 Da, cLogP 1–5, HBD ≤5, HBA ≤10, TPSA ≤140 Å ², RotB ≤10, RingCount ≤6, AromaticRings ≤4, HeavyAtoms 20–50, and FormalCharge between –1 and +1.

Compounds were filtered based on Boltz-2 re-scoring, requiring predicted binding probability >0.8, log_10_(IC50) < –1, and compliance with drug-like physicochemical property criteria. To ensure structural diversity among these high-confidence candidates, fingerprint-based clustering was performed using 1024-bit Morgan fingerprints and the Butina algorithm^25^ with a Tanimoto similarity cutoff of 0.7. PCA was used for two-dimensional visualization of structural diversity. Within each resulting cluster, a representative compound was selected based on the lowest predicted log_10_(IC50) value. This strategy prioritizes potent candidates while minimizing redundancy and ensuring broad scaffold coverage among compounds selected for experimental validation. Known HDAC-related annotations were curated using CAS SciFinder (https://scifinder-n.cas.org/)^27^.

### Model interpretability using Shapley Additive exPlanations (SHAP)

To interpret model predictions, SHapley Additive exPlanations (SHAP)^28^ was applied to quantify the contribution of individual fingerprint bits to predicted binding probabilities. SHAP values were computed for the final BindRankNet model using the GradientExplainer implementation, with a randomly sampled 100,000 PubChem compounds used as the background distribution. SHAP values indicate the direction and magnitude of each feature’s effect on the predicted binding probability.

SHAP analysis was performed on two compound sets: (i) the top 100,000 ranked compounds based on BindRankNet predictions prior to downstream filtering, and (ii) the 528 representatives obtained after physicochemical filtering and structural diversity selection. Global feature importance was estimated by averaging SHAP values across compounds within each set, and agreement between the two sets was quantified using Pearson correlation. For structural interpretation, fingerprint bits with high SHAP values were mapped to molecular substructures using RDKit. Corresponding atom environments were identified via the Morgan fingerprint bitInfo mapping, and the associated atoms and bonds were extracted based on the radius-defined neighborhoods.

### Chemical synthesis of selected compounds

The two novel HDAC11 inhibitors were synthesized by WUXI AppTec. CID 89974511 was prepared in six linear steps (**Supplementary Fig. 5a**). Oxidation of ethyl 2-[4-(hydroxymethyl)piperidin-1-yl]pyrimidine-5-carboxylate with Dess–Martin periodinane gave the 4-formylpiperidine intermediate in 65.5% yield (compound 2). Parallel Boc protection of 2-(6-methyl-1H-indol-3-yl)ethanamine hydrochloride afforded the N-Boc derivative in 99.8% yield, followed by N1-alkylation with 1-iodopropane (99.5% yield) and TFA-mediated Boc deprotection (93.8% yield) to furnish 2-(6-methyl-1-propyl-1H-indol-3-yl)ethanamine. Reductive amination of this amine with the 4-formylpiperidine intermediate using NaBH₃CN provided the ethyl ester in 44.1% yield (compound 8). Final conversion of the ester to the hydroxamic acid with hydroxylamine/NaOH, followed by preparative HPLC, delivered CID 89974511 as the hydrochloride salt in 17.9% yield for the last step.

CID 161174911 was synthesized in four steps (**Supplementary Fig. 5b**). Lewis acid-mediated formylation of 6-methylindan-1-one with diethoxymethoxyethane/BF₃·Et₂O gave the 2-(diethoxymethyl) intermediate in 24.1% yield, which was converted to 5-methyl-1H-indene-2-carbaldehyde via NaBH₄ reduction and acid-mediated dehydration in 32.5% yield (compound 12). Reductive amination of the aldehyde with ethyl 2-[4-(aminomethyl)piperidin-1-yl]pyrimidine-5-carboxylate using NaBH₃CN afforded the ethyl ester intermediate (compound 14) in 54.8% yield. Subsequent hydroxamic acid formation with hydroxylamine/NaOH and preparative HPLC purification furnished CID 161174911 as the hydrochloride salt in 33.7% yield for the last step.

### HDAC11 fluorogenic enzyme inhibition assay

Inhibition of HDAC11 enzyme activity by test compounds was measured using a fluorogenic assay kit (BPS Bioscience, #50687) following the manufacturer’s instructions. The assay was adapted from a 96-well (50 μL) to 384-well format within a reaction volume of 10 μL. Assay performance was verified by enzyme titration in the presence of 1% Dimethyl Sulfoxide (DMSO), confirming a linear increase in signal with increasing HDAC11 concentration, with Trichostatin A included as a control inhibitor in the assay kit.

CID 89974511, CID 161174911, Fimepinostat, and Panobinostat were prepared as 10 mM DMSO stocks and serially diluted (2.5-fold). Compounds (100 nL) were dispensed into assay plates using Echo 650 Acoustic dispenser (Beckman), with DMSO vehicle added to control wells. Reactions were performed with 2 ng/μL HDAC11 and 2 μM substrate in assay buffer supplemented with Bovine Serum Albumin (BSA), with a final DMSO concentration of 1%. Plates were incubated at 37 °C for 30 min, followed by addition of developer solution and incubation for 15 min at room temperature. Fluorescence was measured using a BioTek Neo2 plate reader (Excitation 356/20 nm, Emission 450/20 nm), and background signal from no-enzyme “Blank” values was subtracted. Dose–response curves were fitted using nonlinear regression to estimate IC50 values.

### Statistical analysis

Differences in Boltz-2–predicted binding probability and log₁₀(IC50) distributions were assessed using the two-sided Wilcoxon rank-sum test. Effect sizes were quantified using Cliff’s delta (δ), a nonparametric measure of the probability difference between two distributions. For correlation analyses, Pearson correlation coefficients (r) and Spearman rank correlation coefficients (ρ) were computed. All statistical analyses were performed in R (version 4.3.1) and Python (version 3.8.20).

## Data availability

The PubChem compound library used in this study is publicly accessible at https://pubchem.ncbi.nlm.nih.gov/. The Boltz-2–predicted binding probabilities and log₁₀(IC50) values for the randomly sampled 1.85 million compounds used in this study are available via Zenodo at https://doi.org/10.5281/zenodo.19798185 (ref. ^39^). The list of high-confidence candidate compounds identified by FastBindRank is provided in Supplementary information.

## Code availability

The code for this work is available via GitHub at https://github.com/jwdelta/FastBindRank and via Zenodo at https://doi.org/10.5281/zenodo.19824576 (ref. ^40^).

## Acknowledgements

This work was supported by a Breast Cancer Research Foundation Investigator Award (BCRF-22-133) to L.P. and in part by a National Science Foundation I-Corps award (TI-2449178) to W.H.L. Yale Center for Molecular Discovery is supported in part by an NCI Cancer Center Support Grant (NIH P30 CA016359). This work utilized Echo 650 Acoustic Liquid Handler that was purchased with funding from a National Institutes of Health SIG grant (1S10OD038347). M.M. is supported by a donation from Carmelina Ratto and Carlo Capra.

## Author contributions

J.D. and L.P. conceived and designed the study. J.D. developed the framework and performed the analysis with the help of W.H.L. Y.W. and N.S. provided expertise on HDAC11 biology and small molecule inhibitor discovery. W.H.L. and Q.Y. contributed to the critical evaluation of the study. L.K.G and Y.V.S performed experimental validation of novel inhibitors. L.P. supervised the study. J.D. wrote the original manuscript. Y.W., N.S., M.M., Z.Y., Q.Y., L.K.G., Y.V.S., W.H.L., and L.P. reviewed and edited the manuscript.

## References

1. Cosconati, S. et al. Virtual screening with AutoDock: theory and practice. Expert opinion on drug discovery 5, 597–607 (2010).

2. Eberhardt, J., Santos-Martins, D., Tillack, A. F. & Forli, S. AutoDock Vina 1.2. 0: new docking methods, expanded force field, and python bindings. Journal of chemical information and modeling 61, 3891–3898 (2021).

3. Wang, Y., Wei, Z. & Xi, L. Sfcnn: a novel scoring function based on 3D convolutional neural network for accurate and stable protein–ligand affinity prediction. BMC bioinformatics 23, 222 (2022).

4. Zhang, H. et al. DeepBindBC: A practical deep learning method for identifying native-like protein-ligand complexes in virtual screening. Methods 205, 247–262 (2022).

5. Yang, Z., Zhong, W., Lv, Q., Dong, T. & Yu-Chian Chen, C. Geometric interaction graph neural network for predicting protein–ligand binding affinities from 3d structures (gign). The journal of physical chemistry letters 14, 2020–2033 (2023).

6. Lee, H.-J., Emani, P. S. & Gerstein, M. B. Improved prediction of Ligand–Protein binding affinities by Meta-modeling. Journal of Chemical Information and Modeling 64, 8684–8704 (2024).

7. Wohlwend, J. et al. Boltz-1 democratizing biomolecular interaction modeling. BioRxiv, 2024.2011. 2019.624167 (2025).

8. Passaro, S. et al. Boltz-2: Towards accurate and efficient binding affinity prediction. BioRxiv, (2025).

9. Bender, B. J. et al. A practical guide to large-scale docking. Nature protocols 16, 4799–4832 (2021).

10. Lyu, J. et al. Ultra-large library docking for discovering new chemotypes. Nature 566, 224–229 (2019).

11. Gentile, F. et al. Artificial intelligence–enabled virtual screening of ultra-large chemical libraries with deep docking. Nature Protocols 17, 672–697 (2022).

12. Luttens, A. et al. Rapid traversal of vast chemical space using machine learning-guided docking screens. Nature Computational Science, 1–12 (2025).

13. Graff, D. E., Shakhnovich, E. I. & Coley, C. W. Accelerating high-throughput virtual screening through molecular pool-based active learning. Chemical science 12, 7866–7881 (2021).

14. Marin, E. et al. Regression-based active learning for accessible acceleration of ultra-large library docking. Journal of chemical information and modeling 64, 2612–2623 (2023).

15. Yu, L., He, X., Fang, X., Liu, L. & Liu, J. Deep learning with geometry-enhanced molecular representation for augmentation of large-scale docking-based virtual screening. Journal of Chemical Information and Modeling 63, 6501–6514 (2023).

16. Buciluǎ, C., Caruana, R. & Niculescu-Mizil, A. Model compression. In: Proceedings of the 12th ACM SIGKDD international conference on Knowledge discovery and data mining) (2006).

17. Hinton, G., Vinyals, O. & Dean, J. Distilling the knowledge in a neural network. arXiv preprint arXiv:150302531, (2015).

18. Fukuda, T. et al. Efficient knowledge distillation from an ensemble of teachers. In: Interspeech) (2017).

19. Wang, Y. et al. PubChem: a public information system for analyzing bioactivities of small molecules. Nucleic acids research 37, W623–W633 (2009).

20. Kim, S. et al. PubChem 2023 update. Nucleic acids research 51, D1373–D1380 (2023).

21. Liu, S.-S., Wu, F., Jin, Y.-M., Chang, W.-Q. & Xu, T.-M. HDAC11: a rising star in epigenetics. Biomedicine & pharmacotherapy 131, 110607 (2020).

22. Liu, Y., Tong, X., Hu, W. & Chen, D. HDAC11: A novel target for improved cancer therapy. Biomedicine & Pharmacotherapy 166, 115418 (2023).

23. Khatun, S. et al. Unraveling HDAC11: Epigenetic orchestra in different diseases and structural insights for inhibitor design. Biochemical Pharmacology 225, 116312 (2024).

24. Furui, K. & Ohue, M. Boltzina: Efficient and Accurate Virtual Screening via Docking-Guided Binding Prediction with Boltz-2. arXiv preprint arXiv:250817555, (2025).

25. Butina, D. Unsupervised data base clustering based on daylight’s fingerprint and Tanimoto similarity: A fast and automated way to cluster small and large data sets. Journal of Chemical Information and Computer Sciences 39, 747–750 (1999).

26. Abdi, H. & Williams, L. J. Principal component analysis. Wiley interdisciplinary reviews: computational statistics 2, 433–459 (2010).

27. Gabrielson, S. W. SciFinder. Journal of the Medical Library Association: JMLA 106, 588 (2018).

28. Lundberg, S. M. & Lee, S.-I. A unified approach to interpreting model predictions. Advances in neural information processing systems 30, (2017).

29. Irwin, J. J., Sterling, T., Mysinger, M. M., Bolstad, E. S. & Coleman, R. G. ZINC: a free tool to discover chemistry for biology. Journal of chemical information and modeling 52, 1757–1768 (2012).

30. Liu, X. et al. PRISM: A Structure-Guided Computational Approach for Identifying RNA-Targeting Small Molecules by Integrating 3D Conformational and Chemical Information. European Journal of Medicinal Chemistry, 118795 (2026).

31. Jia, G., Liu, J., Hou, X., Jiang, Y. & Li, X. Biological function and small molecule inhibitors of histone deacetylase 11. European journal of medicinal chemistry 276, 116634 (2024).

32. Gao, L., Cueto, M. A., Asselbergs, F. & Atadja, P. Cloning and functional characterization of HDAC11, a novel member of the human histone deacetylase family. Journal of Biological Chemistry 277, 25748–25755 (2002).

33. Landrum, G. Rdkit documentation. Release 1, 4 (2013).

34. Mirdita, M. et al. ColabFold: making protein folding accessible to all. Nature methods 19, 679–682 (2022).

35. Krivák, R. & Hoksza, D. P2Rank: machine learning based tool for rapid and accurate prediction of ligand binding sites from protein structure. Journal of cheminformatics 10, 39 (2018).

36. Consortium, U. UniProt: a worldwide hub of protein knowledge. Nucleic acids research 47, D506–D515 (2019).

37. Jumper, J. et al. Highly accurate protein structure prediction with AlphaFold. nature 596, 583–589 (2021).

38. Boitreaud, J. et al. Chai-1: Decoding the molecular interactions of life. BioRxiv, (2024).

39. Dai, J. et al. FastBindRank dataset: Boltz-2 predictions of binding probability and log10(IC50) for 1.85 million random PubChem compounds targeting HDAC11. Zenode 10.5281/zenodo.19798185 (2026).

40. Dai, J. et al. FastBindRank software: distillation-based framework for scalable virtual screening across ultra-large chemical libraries. Zenode 10.5281/zenodo.19824576 (2026).

