## Supplementary Figures for "Distillation enables scalable high-fidelity virtual screening across ultra-large chemical libraries"

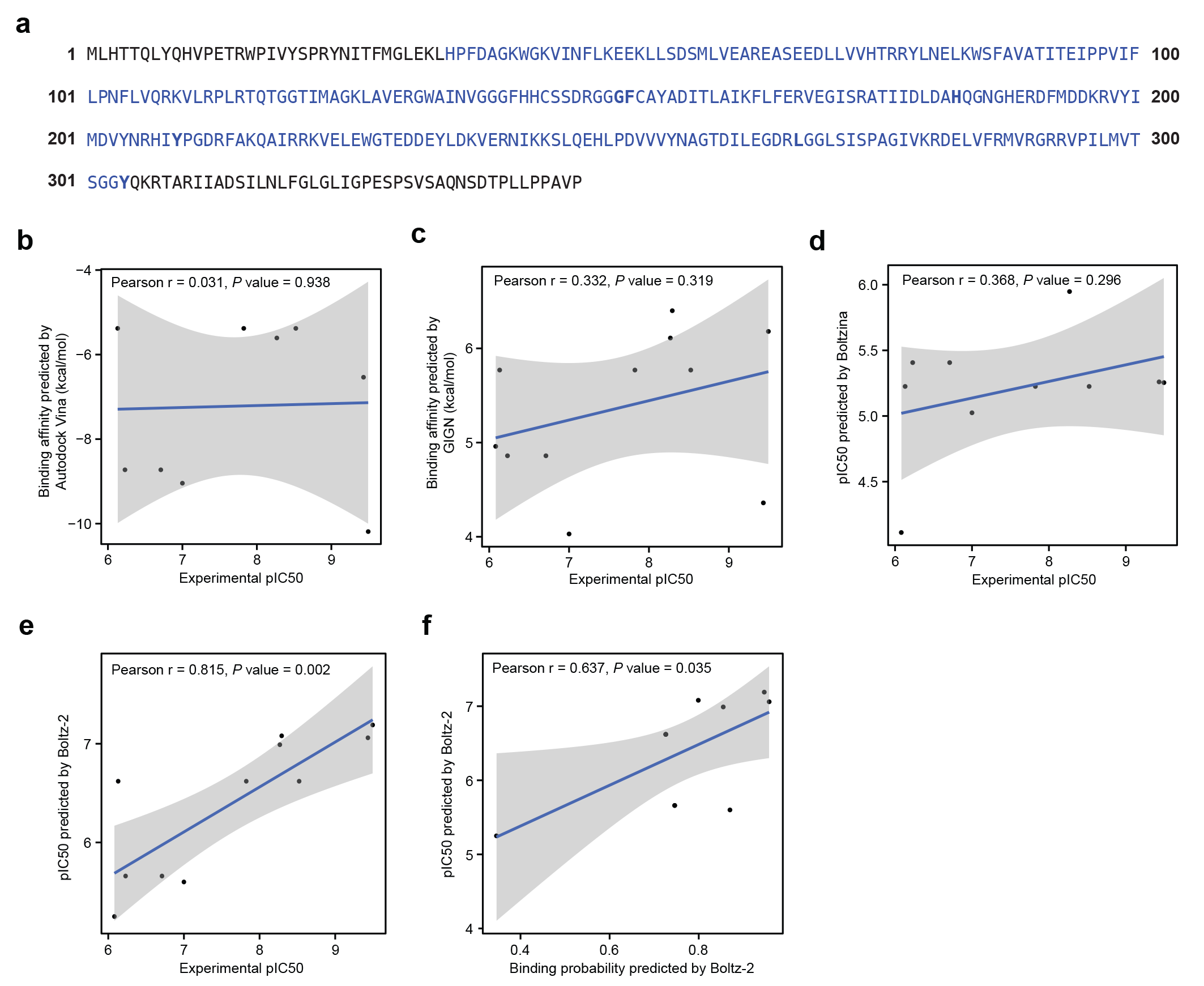

**Supplementary Fig. 1: Sequence annotation of HDAC11 and correlations between predicted and experimental binding affinities for known HDAC11 inhibitors.**

**a,** Amino acid sequence of HDAC11. The catalytic domain is highlighted in blue, and key residues lining the catalytic tunnel are shown in bold. **b–e,** Correlations between predicted binding affinities and experimental pIC50 values measured by enzyme inhibition assays for a curated set of known HDAC11 inhibitors. Predictions are shown for AutoDock Vina (**b**), GIGN (**c**), Boltzina (**d**), and Boltz-2 (**e**). **f,** Correlation between the two Boltz-2 prediction outputs—predicted pIC50 and binding probability—for the same set of known HDAC11 inhibitors. Each point represents one known HDAC11 inhibitor. Blue lines indicate linear regression fits, and shaded regions denote 95% confidence intervals. Pearson correlation coefficients (r) and corresponding *P* values are shown in each panel.

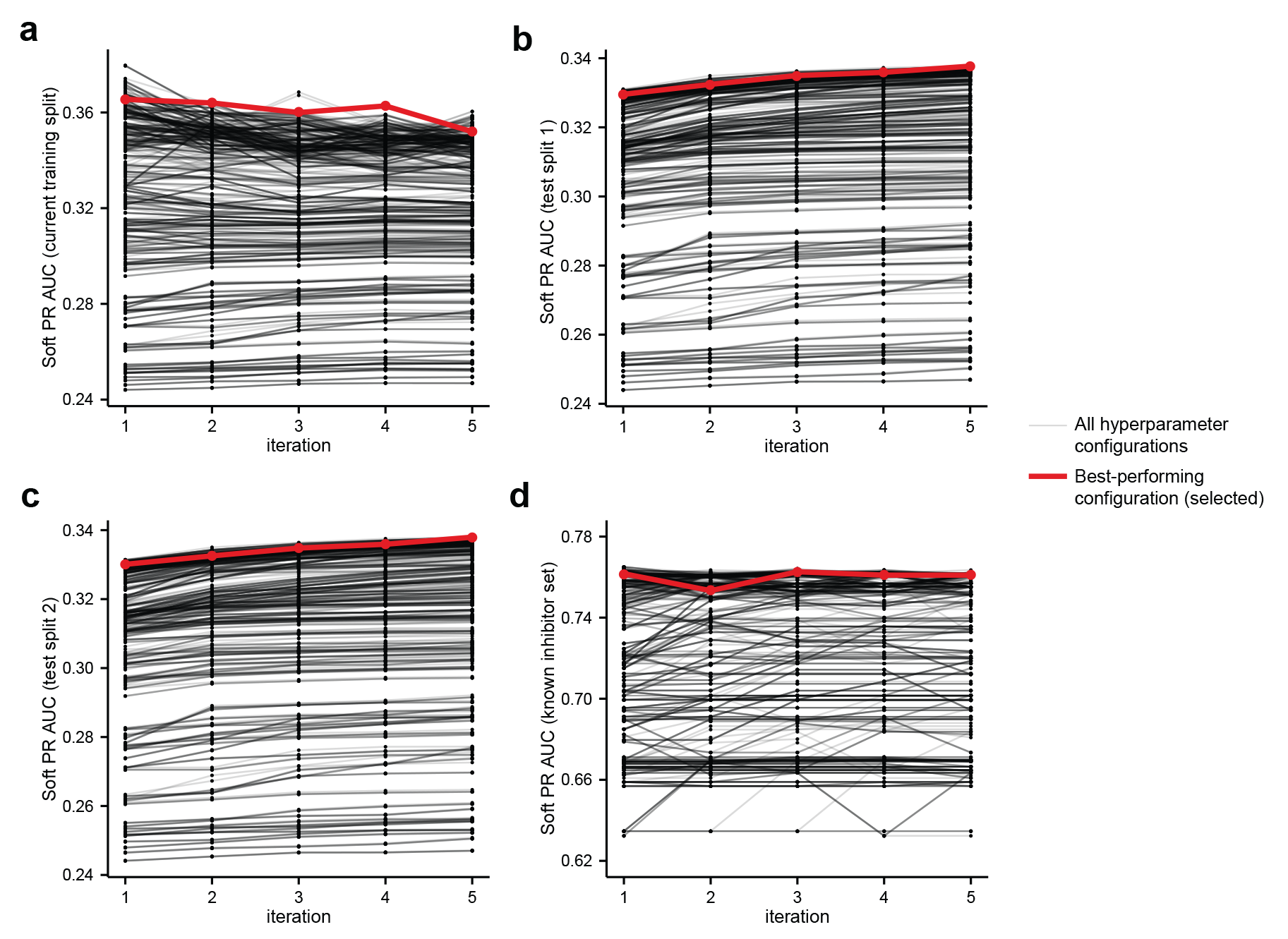

**Supplementary Fig. 2: Performance dynamics of BindRankNet configurations across training iterations.**

**a–d,** Performance of BindRankNet across iterative training rounds for all evaluated hyperparameter configurations, shown on the training splits used at each iteration (**a**), test split 1 (**b**), test split 2 (**c**), and a curated set of known HDAC11 inhibitors (**d**). Each black line represents one hyperparameter configuration, and the red line denotes the best-performing configuration, identified based on soft PR-AUC on the tuning split.

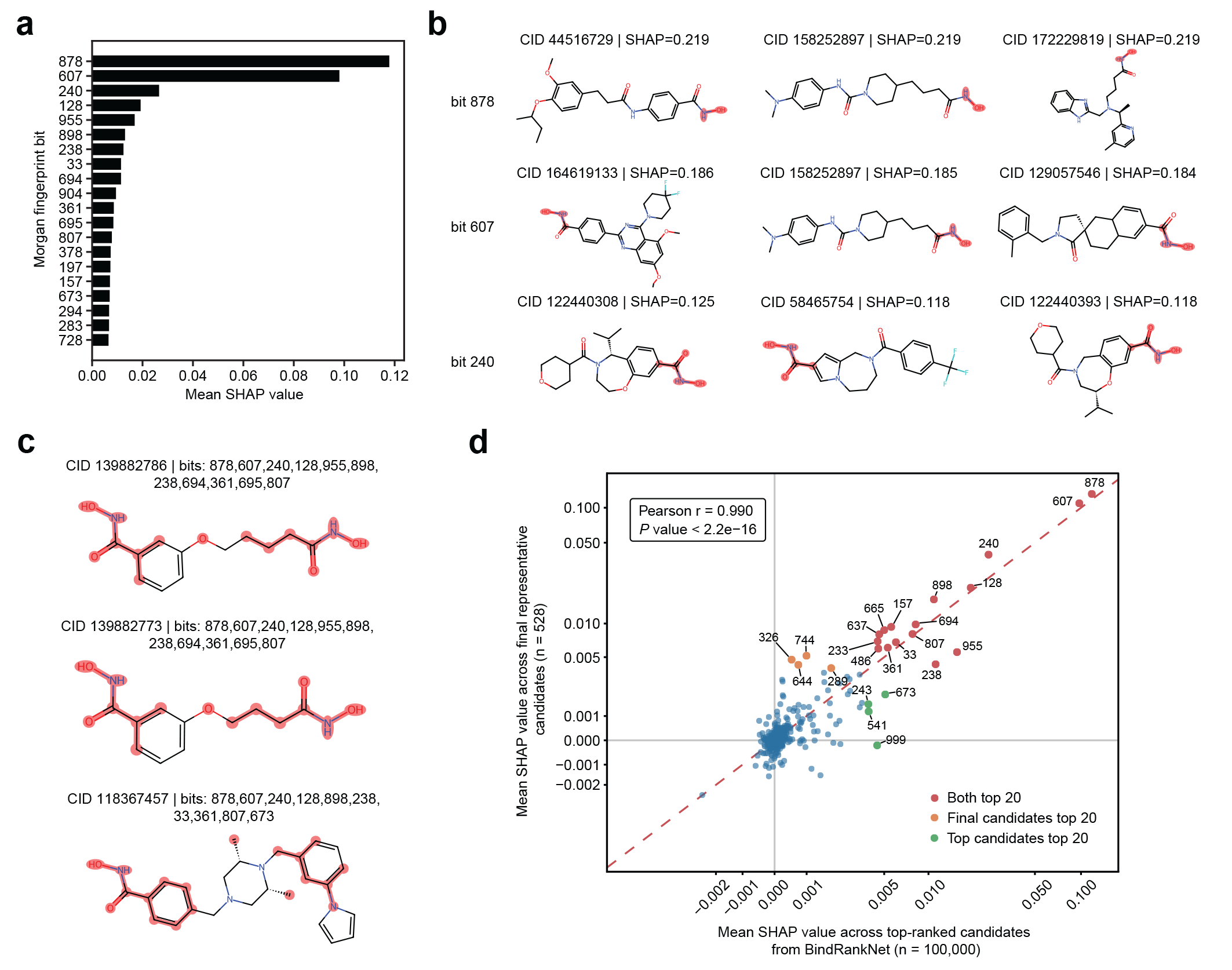

**Supplementary Fig. 3: SHAP-based analysis of structural features associated with predicted HDAC11 binding.**

**a,** Top 20 Morgan fingerprint bits ranked by mean SHAP values across top-ranked candidates predicted by BindRankNet (n = 100,000), highlighting fingerprint bits with the strongest contributions to predicted binding probability. **b,** Representative atom environments (highlighted in red) corresponding to selected high-SHAP fingerprint bits across multiple compounds. Compounds were selected based on the highest SHAP values for each corresponding bit. Each row represents a fingerprint bit, and each column shows its mapped atom environment in different compounds. **c,** Representative candidate compounds with high-SHAP-associated substructures highlighted in red. Compounds were selected based on the number of high-SHAP features (top 20) present in panel a. **d,** Comparison of SHAP values for individual fingerprint bits computed for the top-ranked candidates (n = 100,000) and final representative candidates (n = 528). The axis is displayed using a pseudo-logarithmic scale. Bits were labeled when they rank among the top 20 positive mean SHAP values in either candidate set.

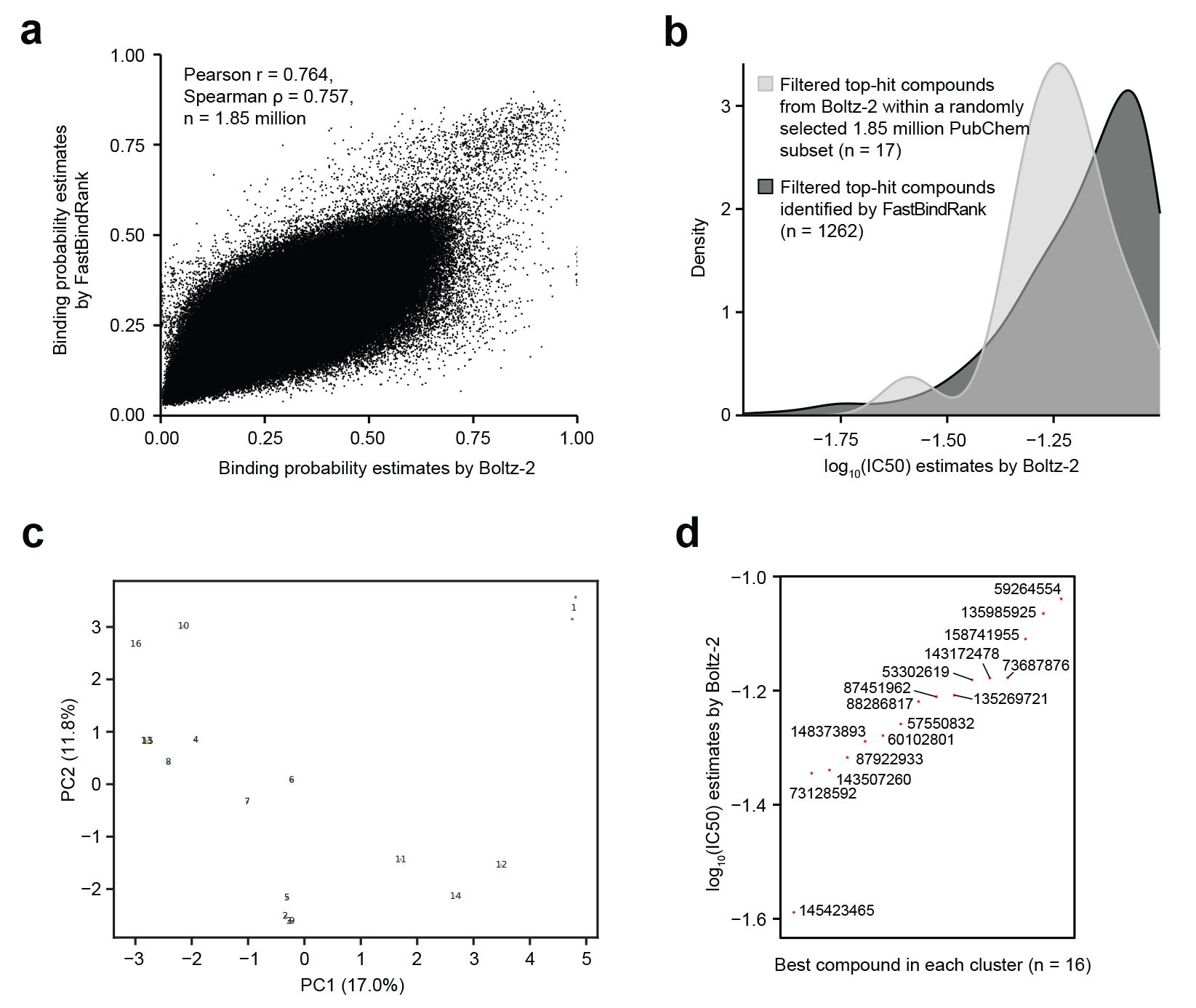

**Supplementary Fig. 4: Subset-level screening behavior.**

**a,** Concordance between binding probabilities predicted by FastBindRank and Boltz-2 for a randomly sampled subset of 1.85 million PubChem compounds. Pearson (r), Spearman (ρ) correlations are shown. **b,** Distribution of Boltz-2–predicted log₁₀(IC50) values for filtered top-hit compounds identified by FastBindRank (n = 1,262; dark grey) compared with filtered top hits from the random subset (n = 17; light grey) using identical filtering criteria including binding probability > 0.8, log₁₀(IC50) < −1, and drug-like molecular properties. **c,** PCA-based visualization of the chemical space of compounds selected from the randomly sampled subset. Numbers denote cluster identifiers. **d,** Rank-ordered Boltz-2–predicted log₁₀(IC50) values for representative compounds selected from each structural cluster (n = 16). These analyses provide context for the subset-level screening behavior used for benchmarking.

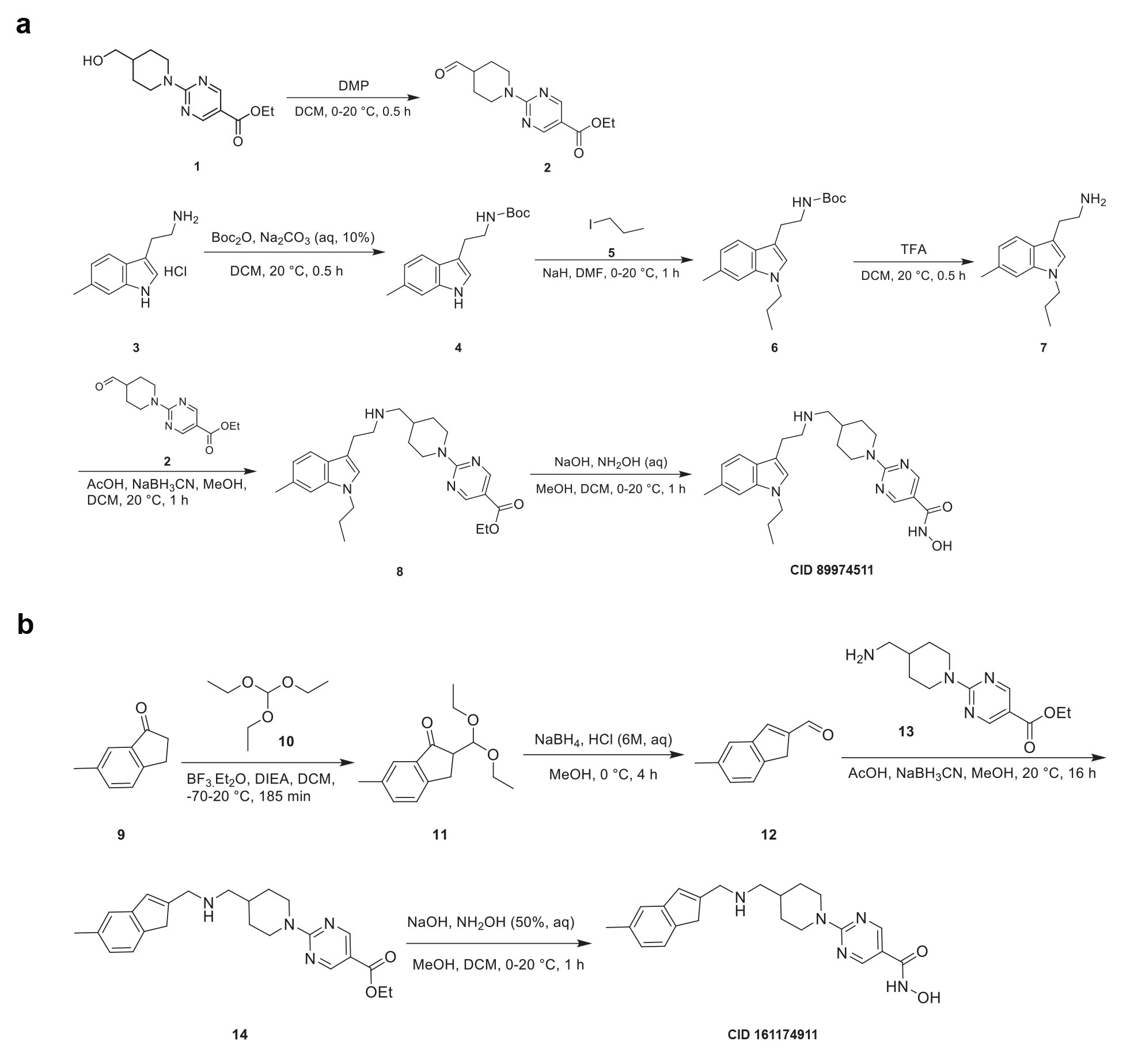

**Supplementary Fig. 5: Synthetic scheme of selected compounds.**

**a,** Synthesis of CID 89974511. **b,** Synthesis of CID 161174911.
